# Detection of Tomato spotted wilt orthotospovirus from crude plant extracts using Reverse Transcriptase- Recombinase Polymerase Amplification in endpoint and real-time

**DOI:** 10.1101/720623

**Authors:** Juan Francisco Iturralde Martinez, Cristina Rosa

**Affiliations:** Department of Plant Pathology and Environmental Microbiology, College of Agricultural Sciences, The Pennsylvania State University, University Park 16802 USA

## Abstract

Virus detection in early stages of infection could prove useful for identification and isolation of foci of inoculum before its spread to the rest of susceptible individuals via vectoring insects. However, the low number of viruses present at the beginning of infection renders their detection and identification difficult and requires the use of highly sensitive laboratory techniques that are often incompatible with a field application.

To obviate this challenge, we designed a Recombinase Polymerase Amplification, a molecular technique that makes millions of copies of a predefined region in the genome, in this case of *Tomato spotted wilt orthotospovirus*. The reaction occurs at 39 ℃ and can be used directly from crude plant extracts without nucleic acid extraction. Notably, positive results can be seen with the naked eye as a flocculus made of newly synthesized DNA and metallic beads.

The objective of the procedure is to create a portable and affordable system that can isolate and identify viruses in the field, from infected plants and suspected insect vectors, and can be used by scientists and extension managers for making informed decisions for viral management. Results can be obtained in situ without the need of sending the samples to a specialized lab.

## Introduction

*Tomato spotted wilt orthotospovirus* (TSWV) causes extensive losses worldwide to various crops and ornamentals ^1^ that can account to as much as 1 billion dollar ^2^. These losses are in part due to the generalist nature of the virus that can infect up to 1000 species of plants, including monocots and dicots ^3^ and has one of the largest host range on record for plant viruses ^4^. Furthermore, TSWV is transmitted in a propagative circulative manner by a dozen polyphagous and hard to control thrips species ^5^.The virus has an ambisense ssRNA genome, comprised of 3 segments named by size small (S), medium (M) and large (L), surrounded by coat proteins and enclosed by a lipidic envelope ^3,6^. Like for other plant viruses, virus and vector exclusion and eradication are the most effective control measures. The use of ultraviolet-reflective mulches, for instance, has been proven to be useful in avoiding primary spread ^7^.

Early detection of TSWV and other plant pathogens is critical for adequate selection and deployment of counter measures ^2^. Common techniques for detection of TSWV include RT-PCR ^8^, immunostrips ^9^, ELISA ^10^ and Next Generation Sequencing ^11^. While molecular assays necessitate expensive lab equipment, can be time consuming and require trained personnel ^12^, antibodies based immunostrips that can be used directly in the field do not offer a high degree of sensitivity, leading to the possibility of false negative and missed virus detection. Losses caused by a lack of detection can be significant, especially for an economically important virus, such as TSWV. Providing a field-based, inexpensive, sensitive and easy-to-use detection assay would allow to test infected material, even before the onset of symptoms, and would result in better disease management.

Recombinase polymerase amplification ^13^ is a nucleic acid sequence based amplification (NASBA) ^14^ that allows amplification, at low constant temperature ^15^. Unlike Loop-mediated Isothermal amplification (LAMP), another isothermal amplification technique which requires a temperature of 65 °C, RPA can perform well at temperatures close to 37 °C, and even ambient temperature, although with lower efficiency ^16^. Because of this, RPA forgoes the need of a thermal cycler and bulky equipment and can be used in the field ^17 18^. Even more conveniently, all reaction components can be lyophilized, allowing room temperature storage and transport under field conditions, and the assay can be performed by untrained personnel ^15^. Mechanistically, RPA exploits a recombination and repair system found in phages ^19^; one of its reagents consist of a nucleoprotein complex (recombinase + primer) that exchange non-template strand and primer, followed by extension by DNA pol, whilst proteins stabilize nascent ssDNA. The end-point product is usually detected using a commercially available amplification detection chamber or lateral flow devices ^20^ which contain proprietary mixes of antibodies that detect FAM and biotin from the amplified products ^21^.

In cases in which there is the need to follow RPA in real time, the design of the probe can be modified to move the FAM molecule closer to the recombination site and quenched in the intact probe. In this way, as the reaction happens, the FAM is cleaved and its concentration can be detected with a fluorometer, as directly proportional to the amount of amplicon being produced ^22^. Advantages of RPA are its readiness and affordability, specificity, sensitivity, speed and easiness of use ^23^ as well as the possibility of multiplexing ^24^. Notably, the sensitivity and specificity of this reaction are comparable and sometimes even superior to that of PCR ^17^. Because of these reasons, RPA is becoming a favorite technique for performing lab-on-a-chip microfluidics analyses that require minimum incubation and can give real-time measurements ^15^.

Binding-flocculation detection of amplicons is a cheaper and equally sensitive approach for detection of RPA products. This techniques exploits the property of aggregation of newly-synthesized nucleic acid and magnetic beads, which form stable floccules under acidic conditions, indicating a positive result ^25^. Interestingly, this same principle can be used for a variety of applications, including the detection of methylation ^26^.

In this work, we present the creation of an RT-RPA assay specific for TSWV detection that can be interpreted with amplification detection chambers, flocculation essays or by real time PCR. We envision that such a test will become a useful addition to the toolkit already in use for TSWV diagnosis.

## Results

### End-point RPA can be visualized in amplification detection chambers

Crude extract from a *Nicotiana benthamiana* (*N. benthamiana*) leaf infected with TSWV was used for RT-RPA reaction and the results of the amplification were visualized in an Amplification detection chamber (Agdia, Elkhart, IN). As seen in Figure 1, both the control (top) and the test line (bottom line) were visible in the chamber where the extract from the infected leaf was loaded (left chamber). This result, indicates a positive amplification, confirming the presence of the dual labelled RT-RPA products and thus of TSWV in the mixture. A negative control, made with the crude extract of a non-infected plant, and one where water was used instead of leaf extract were also subjected to the same procedure. Results in Figure 1 (center and right chambers respectively) show that for both controls only the control line (top line) was visible, indicating a valid test but negative result.

**Figure 1:**
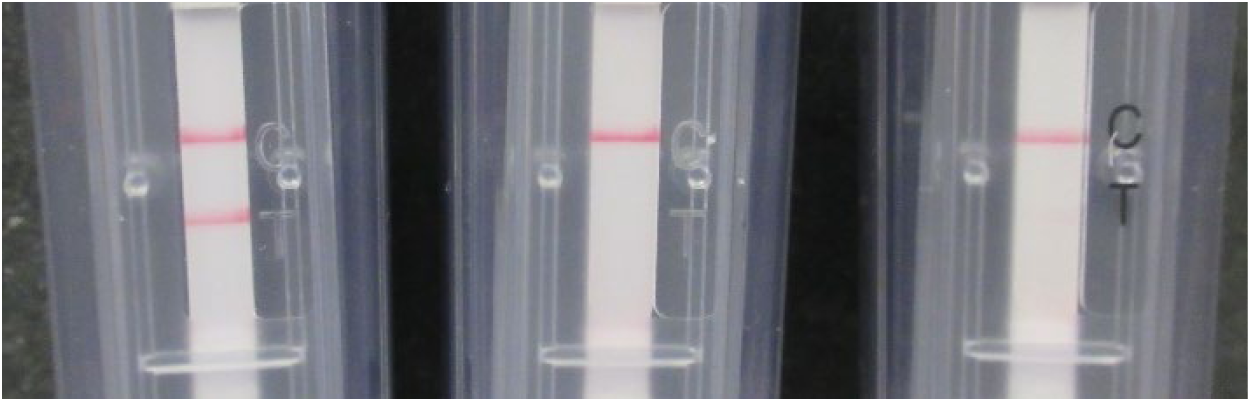
Result of end-point Reverse transcriptase RPA (RT-RPA) in amplification detection chambers (Agdia, Elkhart, IN). Left: positive result for crude extract of a TSWV infected *N. benthamiana* leaf, where both the control (top) and the test line (bottom) are visible. Center and Right: negative results for crude extract of an uninfected leaf and water respectively, where only the control line (top) is visible. NTC1: healthy *N. benthamiana*, NTC2: water.

### End-point RT-RPA can be visualized by flocculation assay

After performing the RT-RPA amplification, the tubes were removed from the thermal block, SPRI (Solid Phase Reversible Immobilization) magnetic beads were added to the amplified fraction, washed with ethanol, followed by the addition of the acetate-based crowding agent. The tube that contained the crude extract of the TSWV infected *N. benthamiana* showed a stable flocculus on the bottom of the tube that did not break, despite flicking the tube repeatedly (Figure 2, left tube). Since the flocculus collects all the beads in the mixture in one small mass, the solution in the tube becomes clear. On the other hand, the vessel containing the non-target controls, did not form a stable flocculus and, due to the even distribution of beads in the mixture, the mixture remained opaque (Figure 2, center and left tube).

**Figure 2:**
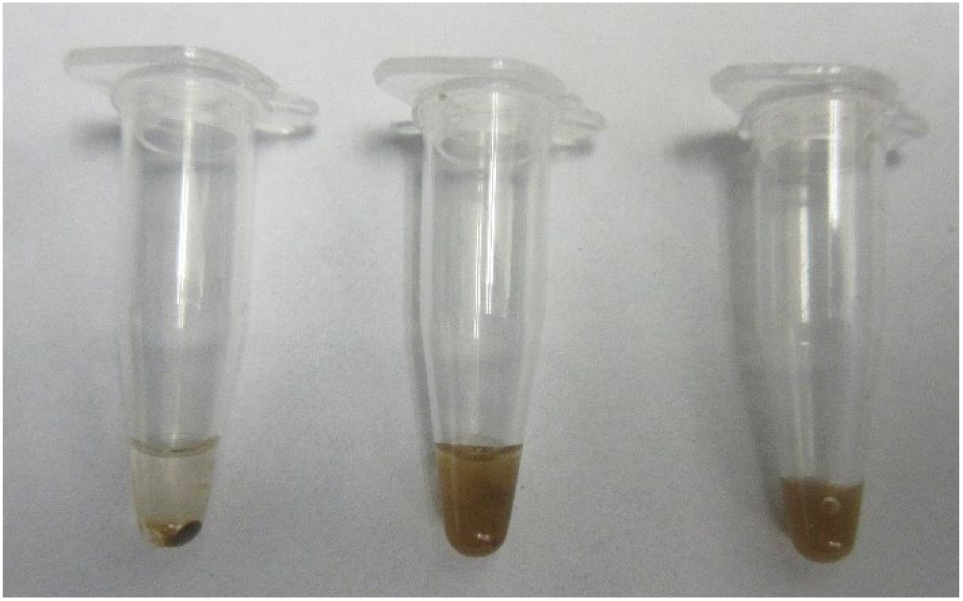
Result of end-point RT-RPA with SPRI beads. Left: positive result, after rehydrating the pellet and the amplification reaction, added SPRI beads react with the amplicon and form a stable flocculus in presence of an acetate-based crowding agent. Center and left: Negative results NTC1: healthy *N. benthamiana*, NTC2: water, the beads are spread throughout the solution making it opaque.

### TSWV RT-RPA assay can detect multiple TSWV isolates in multiple plant species

To ensure that our RT-RPA assay would detect different TSWV isolates and in different plant species, we repeated the RT-RPA reaction for two more TSWV isolates:, TSWV-Chrys5, isolated from Chrysanthemum and maintained in *Nicotiana benthamiana*, TSWV-Tom2, isolated from *Solanum lycopersicum* and maintained in *N. benthamiana*, and TSWV-PA01, isolated from *Capsicum anuum* and maintained in *Emilia sonchifolia* (Margaria and Rosa 2016). All isolates were originally detected via immunostrips (Agdia, Elkhart, IN) and isolated to single lesions to ensure the presence of a single viral species. Our RT-RPA assay was able to detect all the three isolates in all 3 plant species, even if with different sensitivity. Nevertheless, the reaction for TSWV-PA01 was rather weak, when visualized in the amplification detection chambers, but equally positive, when using the flocculation assay.

**Figure 3:**
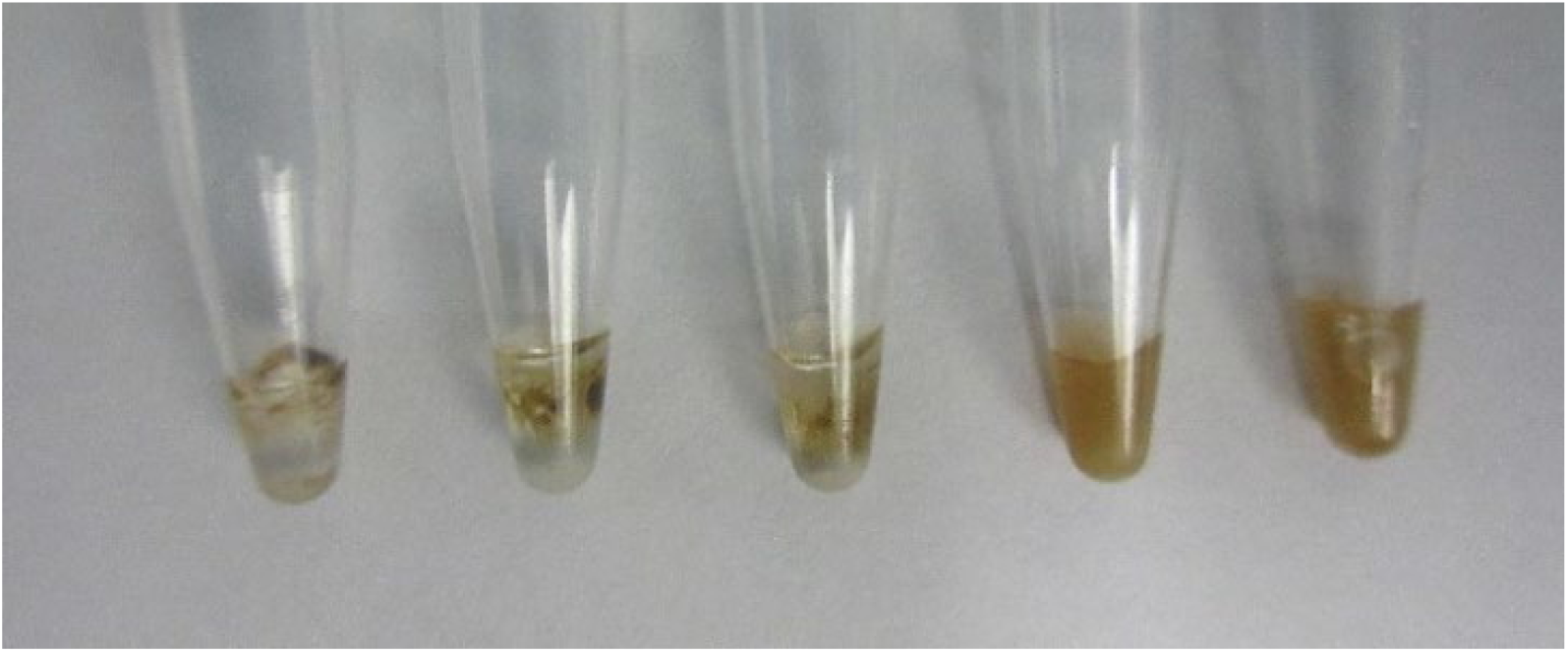
RT-RPA results for three isolates of TSWV revealed with a flocculation assay. TSWV-Chrys5, TSWV-Tom2, TSWV-PA01, NTC1: healthy *N. benthamiana*, NTC2: water.

**Figure 4:**
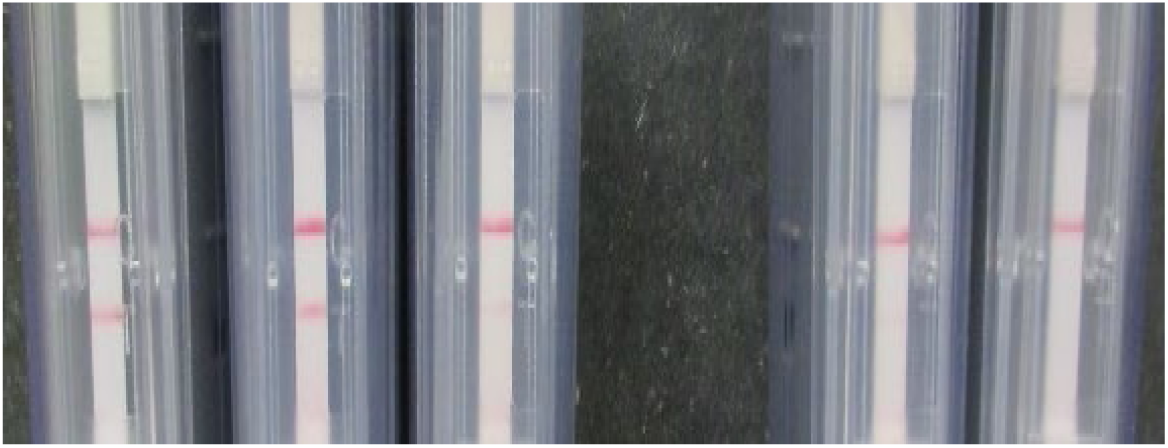
Right: RT-RPA positive result for three isolates of TSWV revealed with an amplification detection chambers. TSWV-Chrys5, TSWV-Tom2, TSWV-PA01 (faint test band), NTC1: healthy *N. benthamiana*, NTC2: water.

### TSWV specific RT-RPA can be visualized by fluorescence in real time

The RT-RPA amplification was followed in real time using the SYBR/FAM channel of a Bio-Rad CFX 96 thermal cycler (Bio-Rad, Hercules, CA). As expected, a positive RT-RPA reaction showed an amplification curve similar to curves typically generated by real-time PCR for the crude extract of a TSWV *N. benthamiana* leaf. The two negative controls, containing both the crude extract of an uninfected leaf and water (non-target controls) did not show an amplification curve.

**Figure 5:**
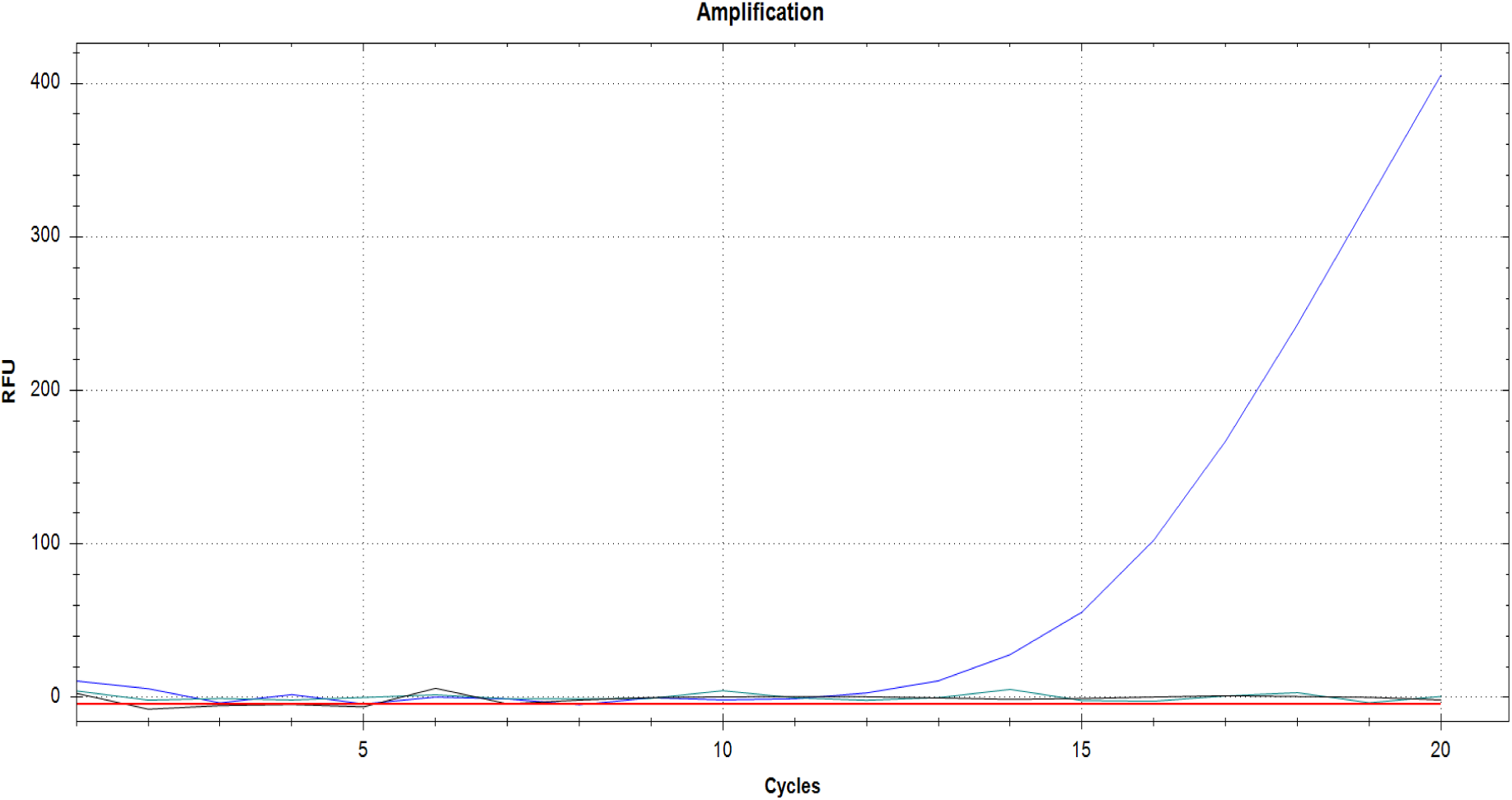
Result of real-time RT-RPA visualized in the SYBR/FAM channel of a Bio-Rad CFX96 real time PCR thermal cycler kept isothermally at 39 °C. Like real time PCR, an amplification curve is visible (blue). The non-target controls (black and green) did not produce an amplification curves in the 20 minutes the reaction took place. Black lines NTC1: healthy *N. benthamiana*, NTC2: water.

### Standard curve

For standard curve analyses, a qRT-PCR product was purified from a completed PCR reaction; then its concentration was measured and used to make serial dilutions. The number of copies of template was calculated using the equation (1)

The standard curve, calculate by making a linear regression of the obtained Cq in the y-axis vs. the log_10_ of the calculated number of copies in the x-axis, is shown in Figure 6.

**Figure 6:**
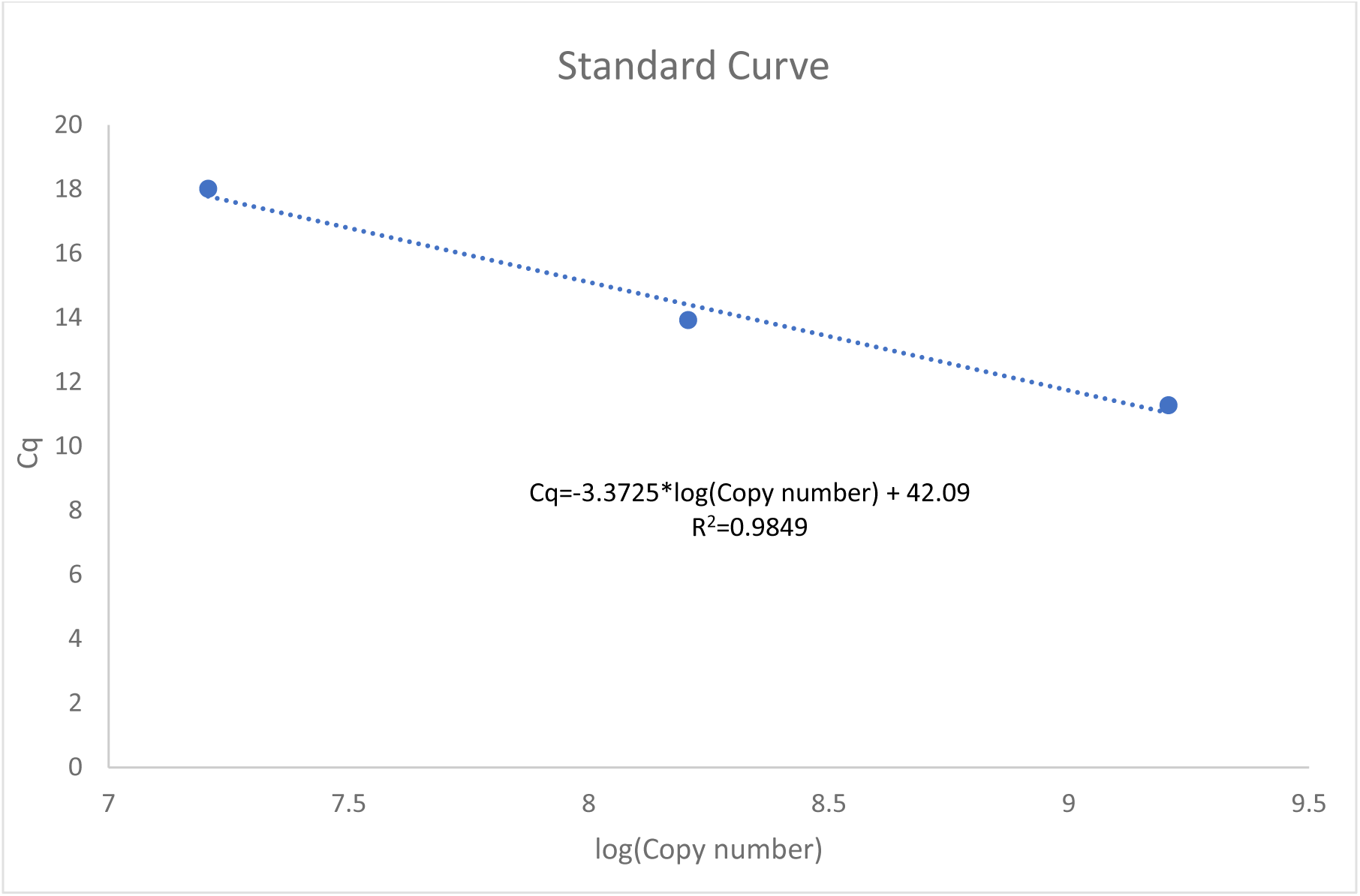
Standard curve computed for absolute quantification. The curve was built by purifying the RT-qPCR amplicon and making 10-fold serial dilutions of it. The obtained Cq for the dilutions was used for a linear regression with the estimated concentrations (measured by nanodrop).

### Quantification of starting material

Fifty microliters of the TSWV infected *N. benthamiana* extract were subjected to RNA extraction, followed by a one-step real time PCR (Figure 7). The calculated Cq for TSWV in 50 microliters of lysate was close to 1.61, which interpolated in the standard curve regression equation (1), shown in Figure 6, gave an approximate of 6.02333E+12 copies of the amplified portion of the TSWV L genomic segment). Since only 2 microliters of this solution were used for the crude extracts and all its dilutions, this gives an estimated initial concentration of 2.40933E+11 genomic segments.

**Figure 7:**
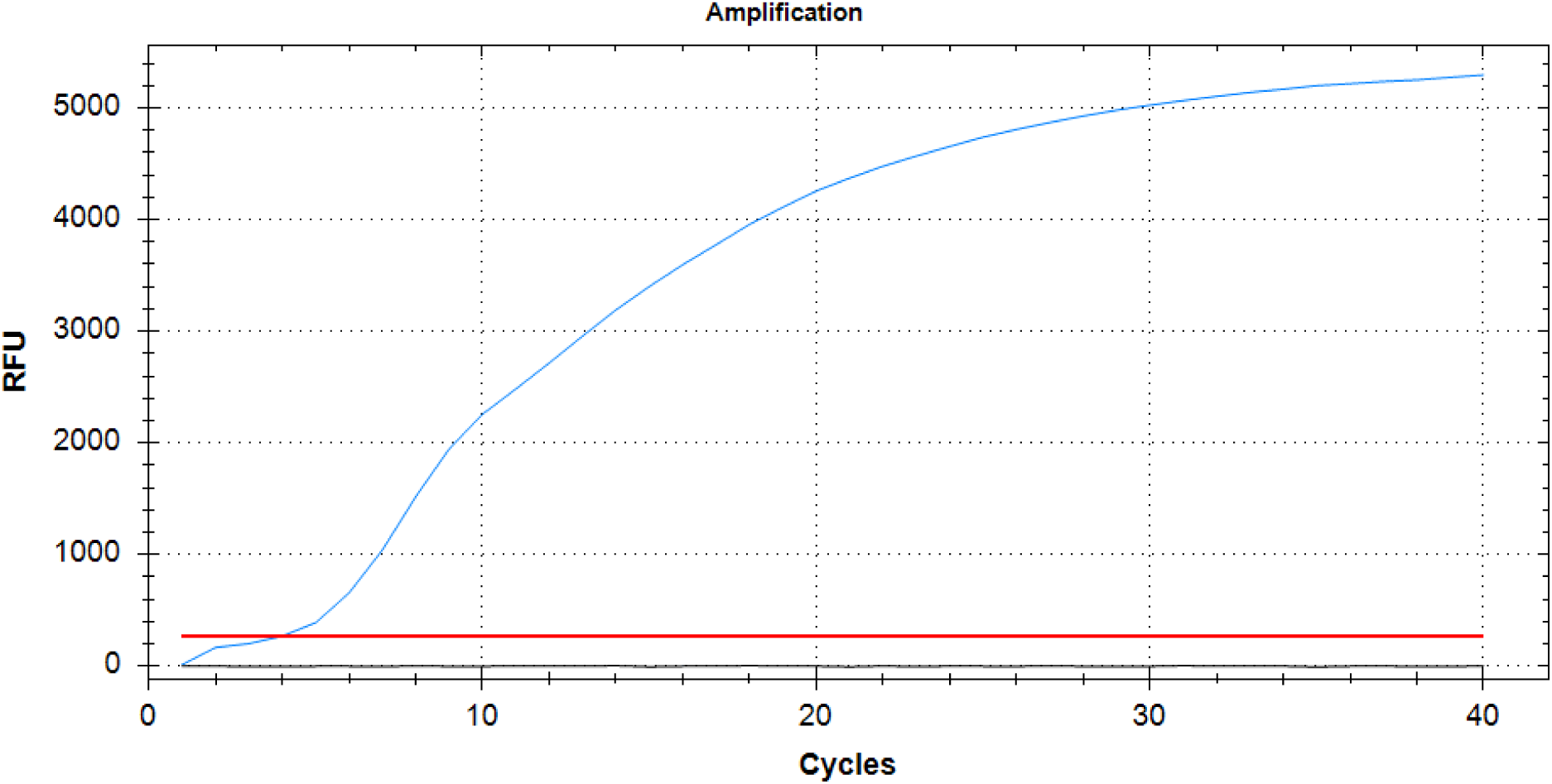
qRT-PCR Amplification curve. RNA was extracted from 50 µl of plant lysate and amplified via one-step qRT-PCR for 40 cycles. The amplification curve is visible with a Cq of 1.61 cycles.

### Sensitivity assay

Serial dilutions of the crude extracts used for the above experiments were subjected to RT-RPA and visualized in a detection chamber (Agdia, Elkhart, IN) .After 20 minutes of incubation, all the dilutions of the crude extract from the TSWV infected plant, from undiluted to 10^−12^ (~1 copy of TSWV genomic segment) showed two bands, indicating a positive result and establishing the theoretical sensitivity of the assay at 1 viral copy. The negative controls showed only the control line (Figure 8).

**Figure 8:**
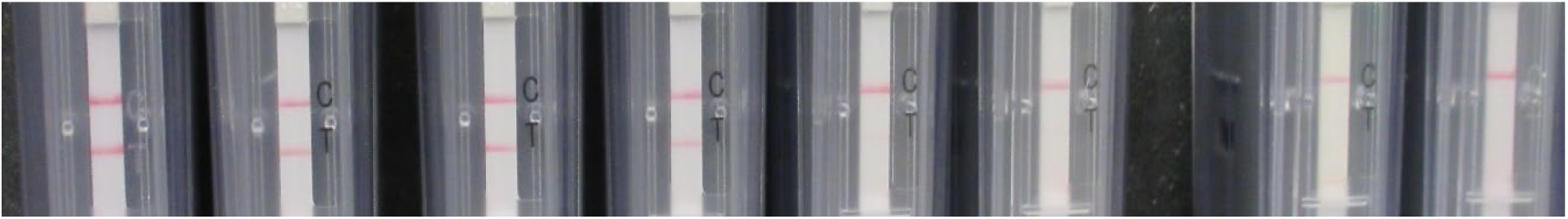
Serial dilutions of a crude extract visualized in amplification detection chambers. From left to right, undiluted sample, 10^−4^ (2^7^ copies), 10^−8^ (2^4^ copies), 10^−12^ (~1 copy) 10^−16^ (out of range), 10^−20^ (out of range) all showing both control and test lines indicating a positive result. To the rightmost side, the two non-target controls NTC1: healthy *N. benthamiana*, NTC2: water were tested, which only showed the control line.

Replicas of the 1:10 dilutions were interpreted with the flocculation assay, showing similar results, all the dilutions, up to 10^−12^ showed a dark precipitate that remained stable even after flicking the tubes repeatedly (Figure 9).

**Figure 9:**
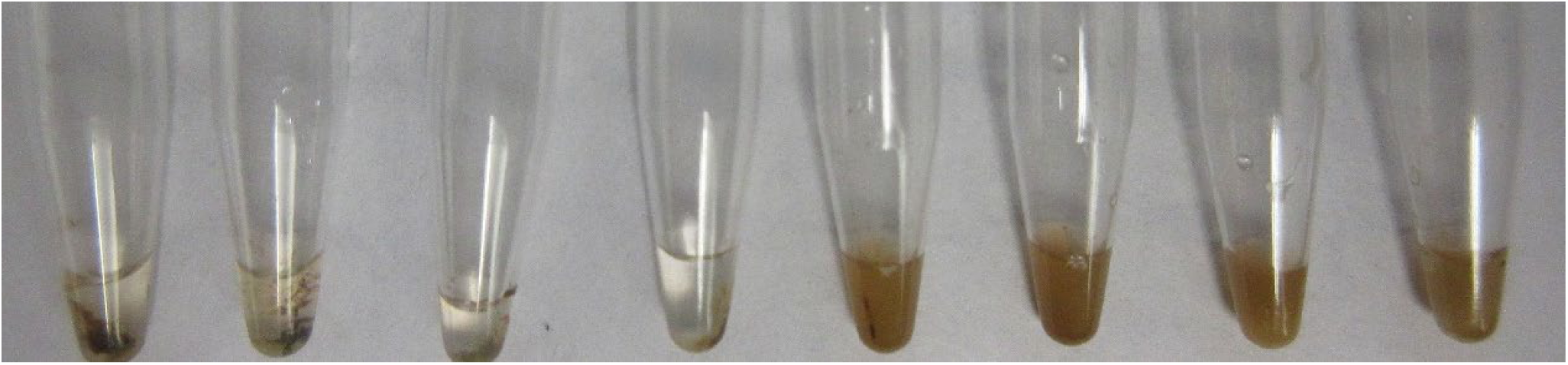
Serial dilutions of a crude extract visualized with SPRI beads. From left to right, undiluted sample, 10^−4^ (2^7^ copies), 10^−8^ (2^4^ copies), 10^−12^ (~1 copy) 10^−16^ (out of range), 10^−20^ (out of range), all showing a flocculus indicating a positive result due to the presence of the amplicon; 10^−16^ and 10^−20^ show no amplification. Last two tubes: NTC1: healthy *N. benthamiana*, NTC2: water.

Finally, the results for real time RT-RPA for some of the above dilutions from undiluted to 10^−20^ in 10^−4^ increments showed differential amplification curves (Figure 10).

**Figure 10:**
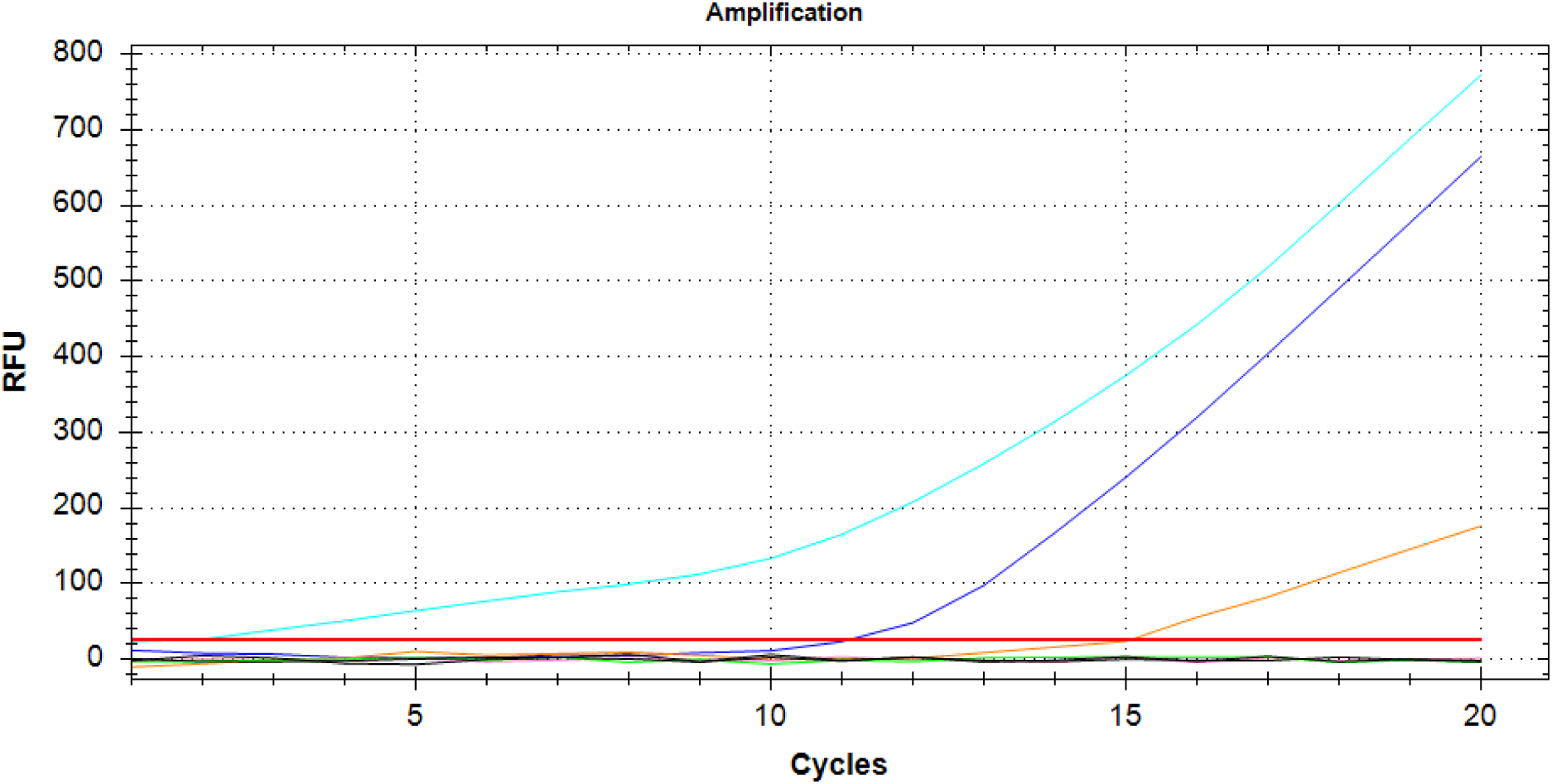
Serial dilutions of a crude extract visualized in the SYBR/FAM channel of a Bio-Rad CFX96 real time PCR thermal cycler kept isothermally at 39 °C From left to right, teal: Crude extract, blue: 10^−4^ (2^7^ copies), orange: 10^−8^ (2^4^ copies), pink: 10^−12^ (~1 copy), green:10^−16^ (out of range), yellow: 10^−20^ (out of range), Black: NTC1: healthy *N. benthamiana*, NTC2: water.

## Discussion and Conclusions

Plant viruses are a threat to food security, because of the huge amount of losses (sometimes as high as 100%, for the case of Tomato spotted wilt virus (TSWV) they cause on crops. As viruses are cellular parasites, no curative measures are available for treating plants, other than prophylactic interventions aimed at controlling their vector and avoid transmission to new hosts ^27^

RT-RPA has been already used for detection of other plant viruses, such as Rose rosette virus ^12^, Plum pox virus ^28^, Little cherry virus-2 ^29^, Yam mosaic virus ^30^ as well as detection at species/subspecies level of Potato virus Y and Wheat dwarf virus ^31^.

The results of the RT-RPA molecular detection technique developed in this study prove that crude extracts of TSWV infected plants can be used directly for sensitive and reliable detection of the virus with a portable but robust technique. In fact, we were able to tailor TSWV specific primers and probes that work with Agdia’s RT-RPA detection chambers and ABM flocculation beads. Our results were confirmed by real time PCR amplification. In the case of real time PCR, it is noteworthy that the appearance of amplification curves obtained was atypical and not linear, suggesting that this assay is not the ideal one to quantify an RT-RPA product, but it should be used more as a qualitative tool for assessing the presence or absence of target. This result is expected, since the design of this assay is suitable for use with a fluorometer and was not intended to be used in real time PCR thermal cyclers. The produced RT-RPA amplicons could also be visualized and characterized using agarose electrophoresis, nonetheless, as other researchers report, the assay can have a carryover of protein that can impose a limitation in the proper migration of the amplified nucleic acid during the electrophoresis if a purification is not performed ^12^.

As seen by Zhang, et al. ^28^ the level of sensitivity of RT-RPA is very high, and allowed us to reach the theoretical detection of one copy of TSWV RNA segment. This unsurpassed sensitivity is intrinsic to RT-RPA, that combines nucleic acid amplification with serological detection.

In conclusion, we were able to design a molecular technique for the rapid and sensitive detection of *Tomato spotted wilt orthotospovirus* that is portable and can be interpreted using three different tools: amplification detection chambers, SPRI beads + crowding agent and fluorometry, using a modified probe. All these techniques can be used interchangeably for a qualitative detection of TSWV and can be scaled up for lab-on-a-chip applications or point-of-care detection of virus, to make quicker informed decisions in resource-limited environments, and without the need of trained personnel.

## Materials and Methods

### Plant rearing and infection

*Nicotiana benthamiana* seeds were germinated in Sunshine #4 aggregate plus soil mix (Sungro, Agawam, MA) for 1 week and the resulting seedlings were transplanted in 4” square pots and reared for 2 weeks. After this, 18 plants were inoculated using around 1g of TSWV-Chrys 5 infected *N. benthamiana* tissue homogenized with mortar and pestle into chilled Paul’s buffer (5 mM diethyldithiocarbamate –DIECA-; 1 mM Disodic Ethylenediaminetetraacetate –EDTA-; 5 mM sodium thioglycolate in 0.05 M phosphate buffer pH 7) ^32,33^ amended with a dash of carborundum and activated charcoal. Other 18 *N. benthamiana* plants were inoculated using TSWV-Tom2. All plants were kept in a growth chamber at 60% relative humidity and a photoperiod of 16/8 hrs day/night for at least 7 days.

Similarly, seeds of *Emilia sonchifolia* were germinated in petri dishes for 1 week and transferred to pots of Sunshine #4 aggregate plus soil mix (Sungro, Agawam, MA) and reared for 2 weeks in a 16/8 hrs day/night photoperiod, relative humidity was not controlled for this case. With the described process, TSWV-PA01 was inoculated and harvested 7 days after.

### Sample homogenization

Leaf tissue was collected 7 days post infection, transferred to an extraction bag (Bioreba, Switzerland) and mixed with 5 ml of phosphate buffered saline (VWR, West Chester, PA), hand homogenized using a hand-held tissue homogenizer (Agdia, Elkhart, IN) and kept at room temperature for the remainder of the experiment. Serial dilutions of this extract were made, when necessary, by mixing 20 µl of extract with 180 µl of nuclease-free water and repeating this process in series for up to 20 times.

### Amplification

Endpoint: AmplifyRP Acceler8 (Agdia, Inc., Elkhart, IN) reaction pellets were rehydrated accordingly to the manufacturer’s instructions. Additionally, iScript Reverse Transcriptase (Bio-Rad, Hercules, CA), was provided externally for converting RNA into cDNA, alongside modified primers and a the Acceler8 probe, both provided by Integrated DNA Technologies (IDT, Coralville, IA).

The probe and primers were designed according to the Assay Design Help Book from Agdia (https://d163axztg8am2h.cloudfront.net/static/doc/b0/4f/4e8d445a51f202f9f57f24a74c8d.pdf). The special considerations to design the probe are here summarized: Less than 50 nt long (42), idSp is the abasic spacer (also known as THF) site that replaced a T, there are 30 spaces between the Fluorophore (FAM) and the abasic site and at least 15 from this site and the 3’ spacer. Our TSWV specific Acceler8 Probe: 5’-/56-FAM/AT TGT ACA AAG GTT TGT TTC GGA ATA AAT CTA GG /idSp/AT TCG CAA CCT AAT CT/3SpC3/-3’ was designed starting at position 3070 to 3163 on the TSWV sequence with accession NC_002050.1 (L segment)

The primers were designed using Primer Blast ^34^ automatic parameters namely, the target was set to TSWV with accession NC_002050.1with the option of searching for other isolates from the Refseq Representative Genome Database by limiting the organism field to *Tomato spotted wilt orthotospovirus* (taxid:1933298), the maximum, optimum and minimum primer sizes were set to 30, 33 and 36 nt respectively, the size of the amplicon was limited from 100 to 230 nt. Finally, a biotin was added to the 5’ end of the reverse primer. The final primer sequences are listed as follows Reverse: 5’-/5BiosG/ATATTGTTATAGAAGGTCCTAATGATT-3’, Forward: 5’-GAATCTATTATACCATTTCTCAATCTCTTAGC −3’.

An abasic site on the middle of a probe is needed for a nuclease to cleave and release the blocking group, allowing the polymerase contained as well in the mixture to extend the probe ^23^. The reverse primer contains a 5’ biotin label and it forms with the probe a dual labeled FAM-biotin amplicon that can be detected with antibodies ^35^.

For performing the reaction, according to the manufacturer’s procedure, dry reaction pellets in 0.2 ml tubes from the AmplifyRP® Discovery Kits (Agdia, Elkhart, IN) were hydrated with: 5.75 μl of rehydration buffer; 0.5 μl of each, primers at 10 μM; 0.25 µl of 10 µM probe, 0.5 µl of iScript Reverse Transcriptase (Bio-Rad, Hercules, CA), were mixed with a 2 microliters of crude plant extract and with a reaction pellet. Following manufacturer’s suggestions, the last 0.5 µl of 280 mM Magnesium acetate (MgOAc) were added to the lid of the microfuge tube and spun down to complete the 10 µl reaction.

Upon closing the vessel, the reaction was maintained at 39 °C for 20 min in a portable thermal block (Agdia, Elkhart, IN), and the results interpreted afterward. For non-target controls, water was used instead of the plant extract, as well as the extract from a healthy plant.

### Revealing of results: Amplification detection chambers

The 0.2 ml tubes were collected from the portable thermal block and inserted in the designated slot in the Amplification detection chambers (Agdia, Elkhart, IN). After this, the tube was smashed into the amplification detection chamber, freeing both the RT-RPA product and the proprietary solution and this device was left undisturbed while the liquids moved upwards through the lateral flow immunostrip for 20 mins.

### Revealing of results: Flocculation essay

For cheaper and in-field visualization methods, we also tried the method of ^36^ for which, two volumes of SPRI (Solid Phase Reversible Immobilization) magnetic beads (Applied Biological Materials, Richmond, BC, Canada) were added to the amplified RT-RPA products, incubated for 5 mins and then vortexed thoroughly for 10 seconds.

The tubes containing the mixture were placed on a magnetic rack and the supernatant was discarded and replaced by 50 µl of 80% ethanol, the mixture was then incubated for 5 min and vortexed 10 seconds again. Finally, the tubes were placed into the magnetic rack again and the ethanol was discarded, the tubes were open to air dry for 5 minutes and 50 µl of crowding solution comprised of 3M sodium acetate and 20% Tween were added.

### Real time

Like in the End point assay, pellets of AmplifyRP Acceler8 (Agdia, Inc., Elkhart, IN) were rehydrated using rehydration buffer, MgOAc, primers, XRT probe and reverse transcriptase. The XRT probe was designed according to the instructions from the Assay design book from Agdia, and synthesized by Integrated DNA Technologies (IDT, Coralville, IA).; the same primers as the end-point essay were used.

The XRT probe has the same sequence as the designed Acceler8 probe but contains a few modifications for being used in real-time, according to the Assay design book. Namely: the FAM was moved from the 5’ end to one side of the abasic site and attached to a T, a black hole quencher was added to the other side of the abasic site and attached to a T nucleotide as well. Probe:5’-ATTGTACAAAGGTTTGTTTCGGAATAAATC/iFluorT/AGG/idSp/AT/iBHQ-1dT/CGCAACCTAATCT/3SpC3/-3’

For this case, to every dried amplification pellet, 5.75 μl of rehydration buffer were added; 0.5 μl of each, primers at 10 μM; 0.25 µl of 10 µl XRT probe, 0.5 µl of iScript Reverse Transcriptase (Bio-Rad, Hercules, CA), were mixed with a 2 microliters of crude plant extract and with a reaction pellet. Following manufacturer’s suggestions, the last 0.5 µl of 280 mM Magnesium acetate (MgOAc) were added to the lid of the microfuge tube and spun down to complete the 10 µl reaction.

For measuring the production of FAM, the vessels were inserted in a Bio-Rad CFX-96 system for real time PCR with a custom-made program that consisted of 20 cycles at 39 °C, with a plate read step every minute. The channel for SYBR/FAM that measures emission at 520 nm was used for collecting fluorescence.

### Standard curve

For standard curve analyses, a qRT-PCR product was purified from a complete PCR reaction using the Zymo DNA Clean & Concentrator™-5 kit (Zymo Research, Irvine, CA) and eluted in 10 μl of nuclease-free water; then its concentration was measured by a Nanodrop 2000 spectrophotometer (Thermo Scientific, Wilmington, DE). Serial dilutions were prepared by adding 10 μl of the original amplicon to 90 μl of nuclease free water for each case. Then, the number of copies of template was calculated using the following formula:

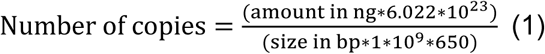

Equation 1. Where 6.022*10^23^ corresponds to the Avogadro’s number for number of molecules per mol, 10^9^ nanograms of exist in one gram and 650 is the average molecular weight in Daltons or g/mol of a pair of bases of DNA (double stranded). The standard curve was made by making a linear regression of the obtained Cq in the y-axis vs. the log_10_ of the calculated number of copies in the x-axis.

### Quantification of starting material

Fifty microliters of *N. benthamiana* lysate were transferred to a 1.5 ml microfuge tube and amended with three volumes of TRIzol LS (Life technologies) and incubated for 3 minutes. Then, a volume of 100% ethanol was added and vortexed for 10 seconds. Finally, Directzol (Zymo Research) kit was used to extract the total RNA, following the manufacturer instructions.

The total RNA was checked in a Nanodrop (ThermoFisher Scientific) spectrophotometer and diluted to 250 ng/µl to be used in real time PCR. The RT-qPCR reaction was performed using a the iTaq™ Universal SYBR® Green One-Step Kit, containing 10 µl of 2X (Bio Rad, Hercules, CA), 0.25 µl of iTaq™ Universal SYBR® Green (Bio-Rad, Hercules, CA), 1X iScript Reverse Transcriptase (Bio-Rad, Hercules, CA), 10 mM of the primers L_TSTW_Fw_Short 5’-GCTCTCCTRTCCACCATCTAC-3’; L_TSTW_Rv_Short 5’-TTGCTGTCAAGGCACACATTTT-3 (Iturralde-Martinez, 2017) designed to amplify a 136 bp amplicon in the TSWV L genomic fragment.

The cycling conditions for the qRT-PCR included a reverse transcription at 50 °C for 10 min, an initial denaturation at 95 °C for 1 min, followed by 40 cycles of denaturation at 95 °C for 10 s, and combined annealing/extending at 60 °C for 30 s, according to the manufacturer’s instructions.

### Sensitivity assay

Serial dilutions of crude extract were made by mixing 10 µl of lysate with 90 µl of nuclease-free water, and then 10 µl of this solution were mixed with 90 µl more and so on (1:10). From each dilution, 2 µl were taken directly and mixed with the master mix for each RT-RPA reaction.

## Acknowledgements

This material is based upon work that is supported by the National Institute of Food and Agriculture, U.S. Department of Agriculture, under award number 2017-38640-26915 through the North Eastern Sustainable Agriculture Research and Education program under subaward number GNE18-176-32231. USDA is an equal opportunity employer and service provider.” C. Rosa is funded by AES # PEN04652.

## References

1 Morsello, S. C. & Kennedy, G. G. Spring temperature and precipitation affect tobacco thrips, Frankliniella fusca, population growth and tomato spotted wilt virus spread within patches of the winter annual weed Stellaria media. Entomologia Experimentalis et Applicata 130, 138–148 (2009).

2 Ohnishi, J. et al. Frankliniella cephalica, a new vector for Tomato spotted wilt virus. Plant disease 90, 685–685 (2006).

3 Pappu, H. Tomato Spotted Wilt Virus. (2008).

4 Hanssen, I. M., Lapidot, M. & Thomma, B. P. Emerging viral diseases of tomato crops. Molecular plant-microbe interactions 23, 539–548 (2010).

5 Webb, S., Tsai, J. & Mitchell, F. in Abstract: Fourth International Symposium on tospoviruses and thrips in floral and vegetable crops. Wageningen, The Netherlands. 67.

6 Adkins, S. Tomato spotted wilt virus—positive steps towards negative success. Molecular Plant Pathology 1, 151–157 (2000).

7 Funderburk, J. et al. Managing thrips in pepper and eggplant. EDIS Document ENY-658, Florida Coop. Ext. Service, Univ. Florida, Gainesville (2011).

8 Roberts, C. A., Dietzgen, R. G., Heelan, L. A. & Maclean, D. J. Real-time RT-PCR fluorescent detection of tomato spotted wilt virus. Journal of Virological Methods 88, 1–8 (2000).

9 Crosslin, J., Mallik, I. & Gudmestad, N. First report of Tomato spotted wilt virus causing potato tuber necrosis in Texas. Plant disease 93, 845–845 (2009).

10 Gonsalves, D. & Trujillo, E. Tomato spotted wilt virus in papaya and detection of the virus by ELISA. Plant Disease 70, 501–506 (1986).

11 Hagen, C. et al. Using small RNA sequences to diagnose, sequence, and investigate the infectivity characteristics of vegetable-infecting viruses. Archives of virology 156, 1209–1216 (2011).

12 Babu, B. et al. A rapid assay for detection of Rose rosette virus using reverse transcription-recombinase polymerase amplification using multiple gene targets. Journal of virological methods 240, 78–84 (2017).

13 Piepenburg, O., Williams, C. H., Stemple, D. L. & Armes, N. A. DNA detection using recombination proteins. PLoS biology 4, e204 (2006).

14 Compton, J. Nucleic acid sequence-based amplification. Nature 350, 91 (1991).

15 Lutz, S. et al. Microfluidic lab-on-a-foil for nucleic acid analysis based on isothermal recombinase polymerase amplification (RPA). Lab on a Chip 10, 887–893 (2010).

16 Crannell, Z. A., Rohrman, B. & Richards-Kortum, R. Equipment-free incubation of recombinase polymerase amplification reactions using body heat. PloS one 9, e112146 (2014).

17 Cabada, M. M. et al. Recombinase polymerase amplification compared to real-time polymerase chain reaction test for the detection of Fasciola hepatica in human stool. The American journal of tropical medicine and hygiene 96, 341–346 (2017).

18 Robertson, B. H. & Nicholson, J. K. New microbiology tools for public health and their implications. Annu. Rev. Public Health. 26, 281–302 (2005).

19 Bianco, P. R., Tracy, R. B. & Kowalczykowski, S. C. DNA strand exchange proteins: a biochemical and physical comparison. Front Biosci 3, D570–D603 (1998).

20 Lillis, L. et al. Factors influencing recombinase polymerase amplification (RPA) assay outcomes at point of care. Molecular and cellular probes 30, 74–78 (2016).

21 Rohrman, B. A. & Richards-Kortum, R. R. A paper and plastic device for performing recombinase polymerase amplification of HIV DNA. Lab on a chip 12, 3082–3088 (2012).

22 TwistDx. (TwistDx Cambridge, UK, 2009).

23 Lobato, I. M. & O’Sullivan, C. K. Recombinase polymerase amplification: Basics, applications and recent advances. Trac Trends in analytical chemistry 98, 19–35 (2018).

24 Crannell, Z. et al. Multiplexed Recombinase Polymerase Amplification Assay To Detect Intestinal Protozoa. Anal Chem 88, 1610–1616, doi:10.1021/acs.analchem.5b03267 (2016).

25 Ng, B. Y., Wee, E. J., West, N. P. & Trau, M. Rapid DNA detection of Mycobacterium tuberculosis-towards single cell sensitivity in point-of-care diagnosis. Scientific reports 5, 15027 (2015).

26 Wee, E. J., Ngo, T. H. & Trau, M. A simple bridging flocculation assay for rapid, sensitive and stringent detection of gene specific DNA methylation. Scientific reports 5, 15028 (2015).

27 Nicaise, V. Crop immunity against viruses: outcomes and future challenges. Frontiers in plant science 5, 660 (2014).

28 Zhang, S. et al. Rapid diagnostic detection of plum pox virus in Prunus plants by isothermal AmplifyRP® using reverse transcription-recombinase polymerase amplification. Journal of virological methods 207, 114–120 (2014).

29 Mekuria, T. A., Zhang, S. & Eastwell, K. C. Rapid and sensitive detection of Little cherry virus 2 using isothermal reverse transcription-recombinase polymerase amplification. Journal of virological methods 205, 24–30 (2014).

30 Silva, G., Bömer, M., Nkere, C., Kumar, P. L. & Seal, S. E. Rapid and specific detection of Yam mosaic virus by reverse-transcription recombinase polymerase amplification. Journal of virological methods 222, 138–144 (2015).

31 Glais, L. & Jacquot, E. in Plant Pathology 207–225 (Springer, 2015).

32 Mautino, G. C., Sacco, D., Ciuffo, M., Turina, M. & Tavella, L. Preliminary evidence of recovery from Tomato spotted wilt virus infection in Frankliniella occidentalis individuals. Annals of applied biology 161, 266–276 (2012).

33 Van Leeuwen, L., Arrieta, I. S., Guiderdone, S. M., Turina, M. & Ciuffo, M. (Google Patents, 2017).

34 Ye, J. et al. Primer-BLAST: a tool to design target-specific primers for polymerase chain reaction. BMC bioinformatics 13, 134 (2012).

35 Daher, R. K., Stewart, G., Boissinot, M. & Bergeron, M. G. Recombinase Polymerase Amplification for Diagnostic Applications. Clin Chem 62, 947–958, doi:10.1373/clinchem.2015.245829 (2016).

36 Koo, K. M., Wee, E. J., Mainwaring, P. N. & Trau, M. A simple, rapid, low-cost technique for naked-eye detection of urine-isolated TMPRSS2: ERG gene fusion RNA. Scientific reports 6, 30722 (2016).

